# ACSL3 is a novel GABARAPL2 interactor that links ufmylation and lipid droplet biogenesis

**DOI:** 10.1101/2020.01.01.892521

**Authors:** Franziska Eck, Manuel Kaulich, Christian Behrends

## Abstract

While studies of ATG genes in knockout models led to an explosion of knowledge about the functions of autophagy components, the exact roles of LC3/GABARAP proteins are still poorly understood. A major drawback for their understanding is that the available interactome data was largely acquired using overexpression systems. To overcome these limitations, we employed CRISPR/Cas9-based genome-editing to generate a panel of cells in which human ATG8 genes were tagged at their natural chromosomal locations with an N-terminal affinity epitope. This cellular resource was exemplarily employed to map endogenous GABARAPL2 protein complexes in response to autophagic modulation using interaction proteomics. This approach identified the ER transmembrane protein and lipid droplet biogenesis factor ACSL3 as a stabilizing GABARAPL2-binding partner. Through this interaction, the GABARAPL2-interacting protein and UFM1-activating enzyme UBA5 becomes anchored at the ER membrane. Functional analysis unveiled ACSL3 and lipid droplet formation as novel regulators of the enigmatic UFM1 conjugation pathway.

## Introduction

From yeast to humans ATG8s are highly conserved proteins. While there is only a single Atg8 in yeast, the human ATG8 (hATG8) family is subdivided into the orthologs microtuble-associated protein 1A/1B light chain 3 (MAP1LC3) including LC3A, LC3B, and LC3C as well as γ-aminobutyric acid receptor-associated protein (GABARAP) including GABARAP, GABARAPL1 and GABARAPL2 (1). All six hATG8 proteins share the same, ubiquitin-like fold although they do not exhibit any sequence homologies with ubiquitin. However, within and between the ATG8 subfamilies, the amino acid sequences show high similarities (2). A major feature of LC3 and GABARAP proteins is their covalent conjugation to the phospholipid phosphatidylethanolamine (PE). This process is initiated by the cysteine proteases ATG4A-D that cleave all hATG8 family members to expose a C-terminal glycine residue and is followed by the activation of LC3s and GABARAPs through the E1-like activating enzyme ATG7. PE-conjugation of hATG8 proteins is subsequently accomplished in a concerted action of the E2-like conjugating enzyme ATG3 and the E3-like ligase scaffold complex ATG12-ATG5-ATG16L1. PE-hATG8 conjugates are reversible through cleavage by ATG4A-D (3).

The best understood function of hATG8s is in macroautophagy (hereafter referred to as autophagy) which is a highly conserved degradation pathway that eliminates defective und unneeded cytosolic material and is rapidly upregulated by environmental stresses such as nutrient deprivation. In the past years, it was shown that autophagy is capable of selectively recognizing and engulfing divers cargo such as aggregated proteins (aggrephagy), pathogens (xenophagy) or mitochondria (mitophagy) with the help of specific receptor proteins (4). Initiation of autophagy leads to the formation of phagophores (also called isolation membranes) from preexisting membrane compartments, such as the ER. Elongation and closure of isolation membranes leads to engulfment of cargo inside double membrane vesicles termed autophagosomes. Fusion of autophagosomes with lysosomes forms autolysosomes in which captured cargo is degraded in bulk by lysosomal hydrolases (5). During this process, GABARAPs and LC3s are associated with the outer and inner membrane of phagophores and regulate membrane expansion (6), cargo receptor recruitment (7), closure of phagophores (8) and the fusion of autophagosomes with lysosomes (9).

Besides autophagy, GABARAPs and LC3s are implicated in a number of other cellular pathways. For example, GABARAP was found as interactor of the GABA receptor and involved in its intracellular transport to the plasma membrane (10, 11), while GABARAPL2 was identified as modulator of Golgi reassembly and intra-Golgi trafficking (12, 13). GABARAPs were also found as essential scaffolds for the ubiquitin ligase CUL3_KBTBD6/KBTBD7_ (14). Among others, LC3s have regulatory functions in RhoA dependent actin cytoskeleton reorganization (15) as well as in the regulation of ER exit sites (ERES) and COPII-dependent ER-to-Golgi transport (16). This high functional diversity of GABARAPs and LC3s implies that these proteins are more than autophagy pathway components and that there are possible other unique functions of individual hATG8 proteins to be unraveled.

So far, interactome and functional analyses of LC3s and GABARAPs were mostly done in cells overexpressing one of the six hATG8 family members (17, 18). This raises the concern that an overexpressed hATG8 protein might take over functions or interactions of one of the other family members due to their high sequential and structural similarity. A lack of isoform specific antibodies further complicates the analysis of distinct functions of hATG8s. To facilitate the study of endogenous GABARAPs and LC3s, it is important to generate alternative resources and tools such as the multiple hATG8 knockout cell lines (9) or the hATG8 family member-specific peptide sensors (19). To circumvent the hATG8 antibody problem, we used CRISPR/Cas9 technology to seamlessly tag hATG8 genes at their natural chromosomal locations. The generated cell lines express N-terminally hemagglutinin (HA)-tagged hATG8 family members at endogenous levels and are a powerful tool to study the functions of individual GABARAPs and LC3s. All created cell lines were tested for their correct sequence and functionality. As a proof of concept, we performed interaction proteomics with the GABARAPL2_endoHA_ cell line and characterized the interaction with the novel binding partner ACSL3.

## Results

### Establishment of cells carrying endogenously HA tagged LC3s and GABARAPs

Complementary to our previously reported LC3C_endoHA_ HeLa cell line (20) we sought to employ CRISPR-mediated gene-editing to generate a panel of cells in which the remaining five hATG8 family members are seamlessly epitope tagged at their natural chromosomal locations. To this end, we directed Cas9 to cleave DNA at the vicinity of the start codon of LC3 and GABARAP genes in order to stimulate microhomology-mediated integration of a sequence encoding for a single HA-tag using a double-stranded DNA donor molecule containing short homology arms (21). Briefly, we designed PCR homology templates in which the blasticidine resistant gene, a P2A sequence and the open reading frame of the HA-tag were flanked by homology arms to the 5’UTRs and first exons of the LC3/GABARAP genes (Supplementary Figure S1A). In parallel, we designed single guide RNAs (sgRNAs) for all hATG8 genes except LC3C and cloned them into pX330 (Addgene 42230), a SpCas9 expressing vector (Supplementary Figure S1A). We then transfected HeLa cells with corresponding pairs of homology template and sgRNA for each LC3/GABARAP gene. After selection with blasticidine, single cell clones were SANGER sequenced to confirm seamless and locus-specific genomic insertion of the HA-tag. While we obtained correct clones for GABARAP, GABARAPL1, GABARAPL2 and LC3B (Supplementary Figure S1B), cells that received the homology template and gRNA for LC3A did not survive the antibiotic selection. We assume that this is due to the lack of LC3A in HeLa cells as it is reported that LC3A expression is suppressed in many tumor cell lines (22). Immunoblot analysis of the sequence-validated clones and the parental cells revealed the presence of the HA-tag in the generated cell lines that corresponded to the size of the tagged LC3/GABARAP protein (Figure 1A, Supplementary Figure S2A-C). Gene specific CRISPR/Cas9-editing was further confirmed by RNAi-mediated depletion of endogenous LC3B or GABARAP proteins in the corresponding HA-tagged hATG8 cell lines (Figure 1B, Supplementary Figure S2D-F). Consistently, confocal microscopy of GABARAPL2_endoHA_ cells showed a substantially decreased HA immunolabeling upon knockdown of GABARAPL2 (Figure 1C). Next, we examined the integrity of the tagged LC3/GABARAP proteins by monitoring their conjugation to PE in response to treatment with small molecule inhibitors which either increase lipidation (Torin1), block autophagosomal degradation (Bafilomycin A1 (BafA1)) or prevent ATG8-PE conjugate formation (ATG7 inhibitor). As expected, GABARAPL2_endoHA_, GABARAP_endoHA_ and LC3B_endoHA_ cell lines showed treatment-specific lipidation levels of the respective tagged hATG8 protein (Figure 1D; Supplementary Figure S2G-I). We also detected lipidated GABARAPL1, though in a manner that was independent from induction or blockage of autophagy (Figure 4C). However, autophagy induction robustly decreased HA-GABARAPL1 protein levels in GABARAPL1_endoHA_ cells while blockage of autophagosomal degradation led to the opposite phenotype (Supplementary Figure S2H). Next, we analyzed the subcellular distribution of one of the HA-tagged hATG8 proteins (i.e. GABARAPL2) in basal and autophagy-modulating conditions using confocal microscopy. In GABARAPL2_endoHA_ cells, HA-GABARAPL2 was indeed found to colocalize with the autophagosomal and -lysosomal markers p62, LC3B and LAMP1 and this colocalization increased upon combination treatment with Torin1 and BafA1 (Figure 1E, Supplementary Figure S2J,K). Together, we successfully engineered cell lines to carry epitope tagged hATG8 family members which retain their functionality.

**Figure 1.**
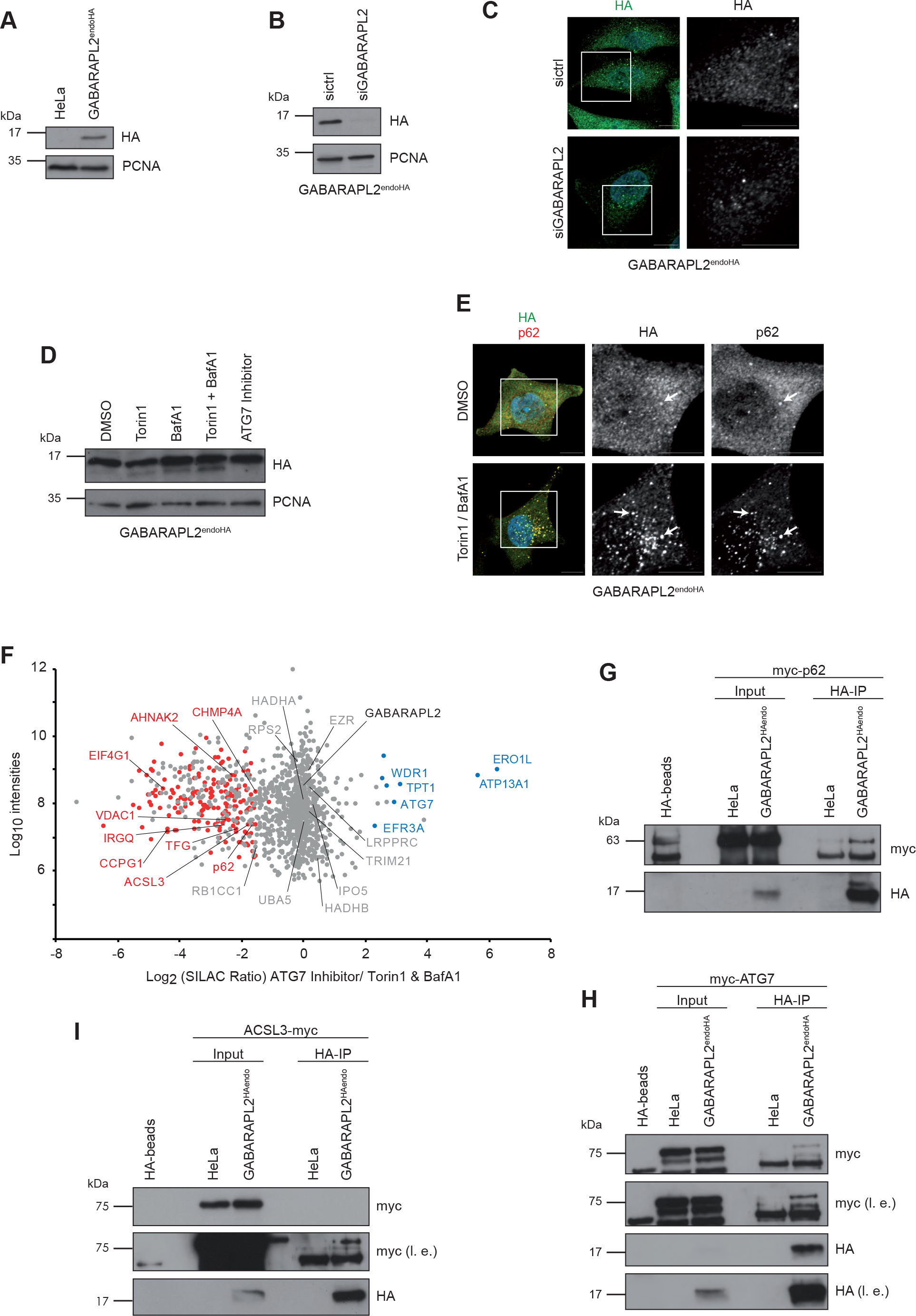
Interaction analysis of endogenously tagged hATG8 proteins. ***A*,** GABARAPL2__endoHA__ and parental HeLa cell lysates were analyzed by immunoblotting using anti-HA and -PCNA antibodies. The latter was used as loading control. ***B,C***, GABARAPL2_endoHA_ cells were reversely transfected for 72 hrs with non-targeting (sicrtl) or GABARAPL2 siRNA followed by lysis and immunoblot analysis (***B***) or fixation and immunolabeling (***C***) using an anti-HA antibody. Scale bar: 10 µm. ***D,E,*** GABARAPL2_endoHA_ cells were treated as indicated and subjected to lysis and immunoblotting (***D***) or fixation and immunolabeling (***E***) using anti-HA and -p62 antibodies. Scale bar: 10 µm. Arrowheads indicate colocalization events. ***F,*** Scatterplot represents interaction proteomics of SILAC labeled GABARAPL2_endoHA_ cells differentially treated with Torin1 and BafA1 (light) or ATG7 inhibitor (heavy). Significantly enriched proteins upon Torin1 and BafA1 combination treatment or ATG7 inhibition are highlighted in red and blue, respectively. Proteins in grey are unchanged. ***G-I,*** Immunoblot analysis of anti-HA immunoprecipitates from lysates derived from parental HeLa and GABARAPL2_endoHA_ cells transiently transfected for 48 hrs with myc-tagged ATG7 (***G***), p62 (***H***) or ACSL3 (***I***).

### Mapping the endogenous GABARAPL2 interactome

Next, we selected GABARAPL2_endoHA_ cells for a proof-of-principle immunoprecipitation (IP) followed by mass spectrometric (MS) analysis to identify new binding partner candidates of a hATG8 family member at endogenous levels. To distinguish between candidates that bind preferentially to PE-conjugated versus unconjugated GABARAPL2 we treated stable isotope labeling with amino acids in cell culture (SILAC)-labeled GABARAPL2_endoHA_ cells with Torin1 and BafA1 (light) or ATG7 inhibitor (heavy). Equal amounts of heavy and light SILAC cells were mixed, lysed and subjected to HA-IP. Immune complexes were eluted and size separated by gel electrophoresis followed by in-gel tryptic digest, peptide extraction and desalting prior to analysis by liquid chromatography tandem MS. SILAC labeled parental HeLa cells differentially treated with Torin1/BafA1 or ATG7 inhibitor served as negative controls. In duplicate experiments, we identified a total of 168 proteins whose abundances in GABARAPL2 immunoprecipitates were altered by at least 2.8-fold (log2 SILAC ratio ≥1.5 or ≤-1.5) in response to modulation of the hATG8 conjugation status (Figure 1F). Among these regulated proteins were well-characterized hATG8 binding proteins such as ATG7, CCPG1 and SQSTM1 (also known as p62) as well as several candidate interaction proteins previously found in large-scale screening efforts such as the mitochondrial outer membrane protein VDAC1, the nucleoprotein AHNAK2, the translation initiation factor EIF4G1 and the small GTPase IRGQ (23, 24) (Figure 1F). In addition, a number of known hATG8 interactors including UBA5, HADHA, HADHB, RB1CC1, TRIM21 and IPO5 was found to bind GABARAPL2 independent of its lipidation status since these proteins did not display substantial changes in their SILAC ratios.

### ACSL3 is a novel binding partner of GABARAPL2

Since functional annotation analysis using DAVID revealed ‘fatty acid metabolism’ as a term previously not associated with LC3/GABARAP-interacting proteins (Supplementary Figure S3A), we focused on the proteins found in this category. In particular, the long-chain-fatty-acid-CoA ligase 3 (ACLS3) attracted our attention as it was the only ER-localized transmembrane protein among these candidates. To validate ACSL3 as novel GABARAPL2 interacting protein, we transiently transfected parental and GABARAPL2_endoHA_ cells with C-terminally myc-tagged ACSL3 followed by HA- or myc-IP and immunoblotting. Transfection with N-terminally tagged ATG7 or p62 served as positive controls. Indeed, ACSL3-myc as well as myc-ATG7 and -p62 associated with endogenous GABARAPL2 (Figure 1G-I, Supplementary Figure S3B). Thus, these results indicate that our hATG8_endoHA_ cells are valuable tools to examine the LC3 and GABARAP interactome at endogenous levels and to identify novel binding partners such as ACSL3.

### ACSL3 recruits GABARAPL2 to the ER membrane

ACSL3 is one of five acyl-CoA synthetases and catalysis the conjugation of CoA to long chain fatty acids to form acyl-CoA (25). Besides ACSL3 was found to regulate the formation, the size and the copy number of lipid droplets (26, 27). Consistent with its cellular role, ACSL3 is located with its N-terminal transmembrane helix region inserted midway into the lipid bilayer of the ER membrane or integrated into the monolayer of lipid droplets (LD) while its C-terminal part encompassing the AMP-binding domain is facing to the cytoplasm (28–30). To further validate the GABARAPL2-ACSL3 interaction, we sought to examine the subcellular localization of both proteins by confocal microscopy. However, as there were no suitable antibodies for immunofluorescence staining of endogenous ACSL3, we gene-edited GABARAPL2_endoHA_ cells to express ACSL3 tagged at its C-terminus with NeonGreen (Supplementary Figure S1A,C). Immunoblot analysis of these newly established GABARAPL2_endoHA_/ACSL3_endoNeonGreen_ cells in comparison with GABARAPL2_endoHA_ and parental Hela cells transfected with TOMM20-NeonGreen confirmed the correct size of the ACSL3-NeonGreen fusion (Figure 2A). Furthermore, colocalization of ACSL3-NeonGreen with the ER-membrane localized chaperone Calnexin demonstrated that the NeonGreen tag did not alter the presence of ACSL3 at the ER (Figure 2B). As ACSL3 is essential for LD formation, we tested whether the ACSL3-NeonGreen chimera is fully functional. Thereto, GABARAPL2_endoHA_/ACSL3_endoNeonGreen_ cells were treated with oleic acid to induce LD formation or EtOH as control prior to fixation and immunolabeling of phospholipids and neutral lipids. Confocal microscopy showed a clear colocalization of ACSL3 with phospholipids and neutral lipids in control cells while ACSL3 redistributed in the phospholipid monolayer of LDs when cells were treated with oleic acid for 24 hrs (Figure 2C). Next, we microscopically analyzed fixed and HA-immunolabeled GABARAPL2_endoHA_/ACSL3_endoNeonGreen_ cells that were grown in the absence and presence of Torin1 and BafA1 or ATG7 inhibitor. Consistent with our IPs, these experiments revealed a partial co-localization of endogenous GABARAPL2 and ACSL3 (Figure 2D). Together, these results show that NeonGreen tagged ACSL3 is correctly localized at the ER membrane, integrates into the monolayer of LDs upon free fatty acid treatment and associates with GABARAPL2 at the ER.

**Figure 2.**
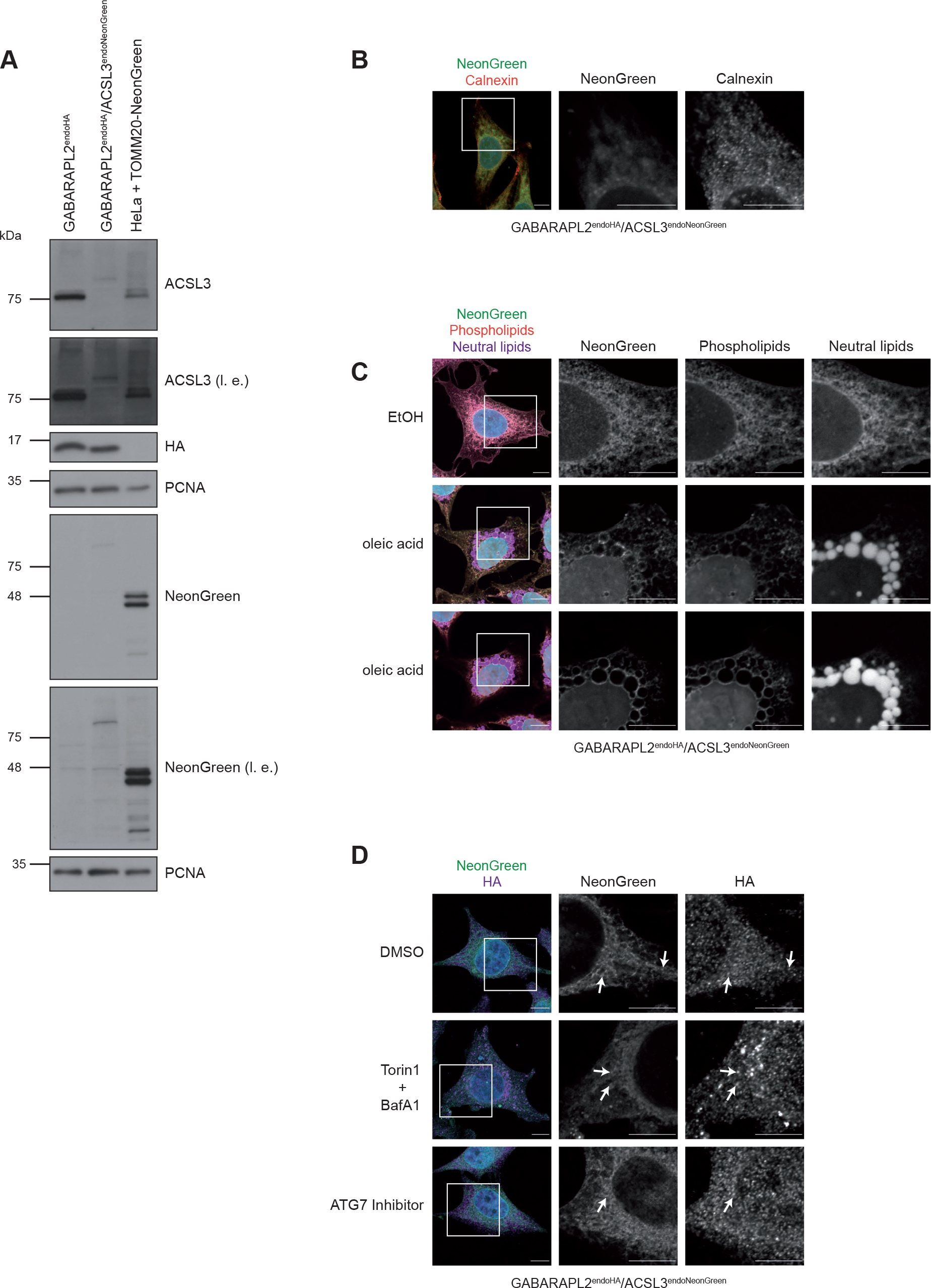
Validation of GABARAPL2_endoHA_/ACSL3_endoNeonGreen_ cells. ***A,*** GABARAPL2_endoHA_ and GABARAPL2_endoHA_/ACSL3_endoNeonGreen_ cells as well as parental HeLa cells transiently transfected with TOMM20-NeonGreen were lysed and analyzed by immunoblotting with indicated antibodies. ***B,*** Fixed GABARAPL2_endoHA_/ACSL3_endoNeonGreen_ cells were immunostained with an anti-calnexin antibody. Scale bar: 10 µm. ***C,*** GABARAPL2_endoHA_/ACSL3_endoNeonGreen_ cells were treated with oleic acid or EtOH (control) for 24 hrs followed by fixation and immunolabeling of phospholipids and neutral lipids. Scale bar: 10 µm. Two confocal planes are shown for oleic acid treatment. ***D,*** GABARAPL2_endoHA_/ACSL3_endoNeonGreen_ cells treated with Torin1 and BafA1 or ATG7 inhibitor were fixed and immunolabeled with an anti-HA antibody. Scale bar: 10 µm. Arrowheads indicate HA-NeonGreen colocalization events.

### GABARAPL2 is stabilized by ACSL3

Since GABARAPL2 is involved in autophagic cargo engulfment (31), we tested whether ACSL3 is an autophagy substrate or serves as selective autophagy receptor. However, stimulation of GABARAPL2_endoHA_ cells with Torin1, BafA1, a combination of both or with ATG7 inhibitor showed that ACSL3 protein levels did not change upon autophagy induction or blockage (Supplementary Figure S3D). Likewise, depletion of GABARAPL2 had no effects on ACSL3 abundance (Supplementary Figure S3E). Thus, these results indicate that ACSL3 is neither a substrate nor a receptor of autophagy under these conditions. Next, we examined the effects of ACSL3 knockdown on GABARAPL2. Treatment of GABARAPL2_endoHA_ cells with two different ACSL3 siRNAs showed a significant decrease of GABARAPL2 protein levels (Figure 3A). To rule out that this phenotype is due to a global perturbation of the ER, we probed for the integrity of this organelle in cells depleted of ACSL3 using immunolabeling with Calnexin and the ER exit site marker SEC13. However, neither the meshwork appearance nor the exist sites of the ER showed any overt alterations (Supplementary Figure S3F,G). Given the high structural and functional similarity between LC3 and GABARAP family members we addressed whether ACSL3 depletion likewise impacts on the protein abundance of the other hATG8 family members. Unexpectedly, ACSL3 knockdown experiments in GABARAP_endoHA_, GABARAPL1_endoHA_ and LC3B_endoHA_ cells did not show any significant reduction in the respective HA-tagged hATG8 proteins (Figure 3B-D). In contrast, we found that LC3B protein levels significantly increased upon ACSL3 depletion (Figure 3D), suggesting that reduced GABARAPL2 levels might be compensated by increased expression of LC3B. Intriguingly, we observed that GABARAPL2 protein levels are restored in RNAi-treated GABARAPL2_endoHA_ cells treated with BafA1 but not with the proteasome inhibitor Bortezomib (Btz) (Figure 3E).

**Figure 3.**
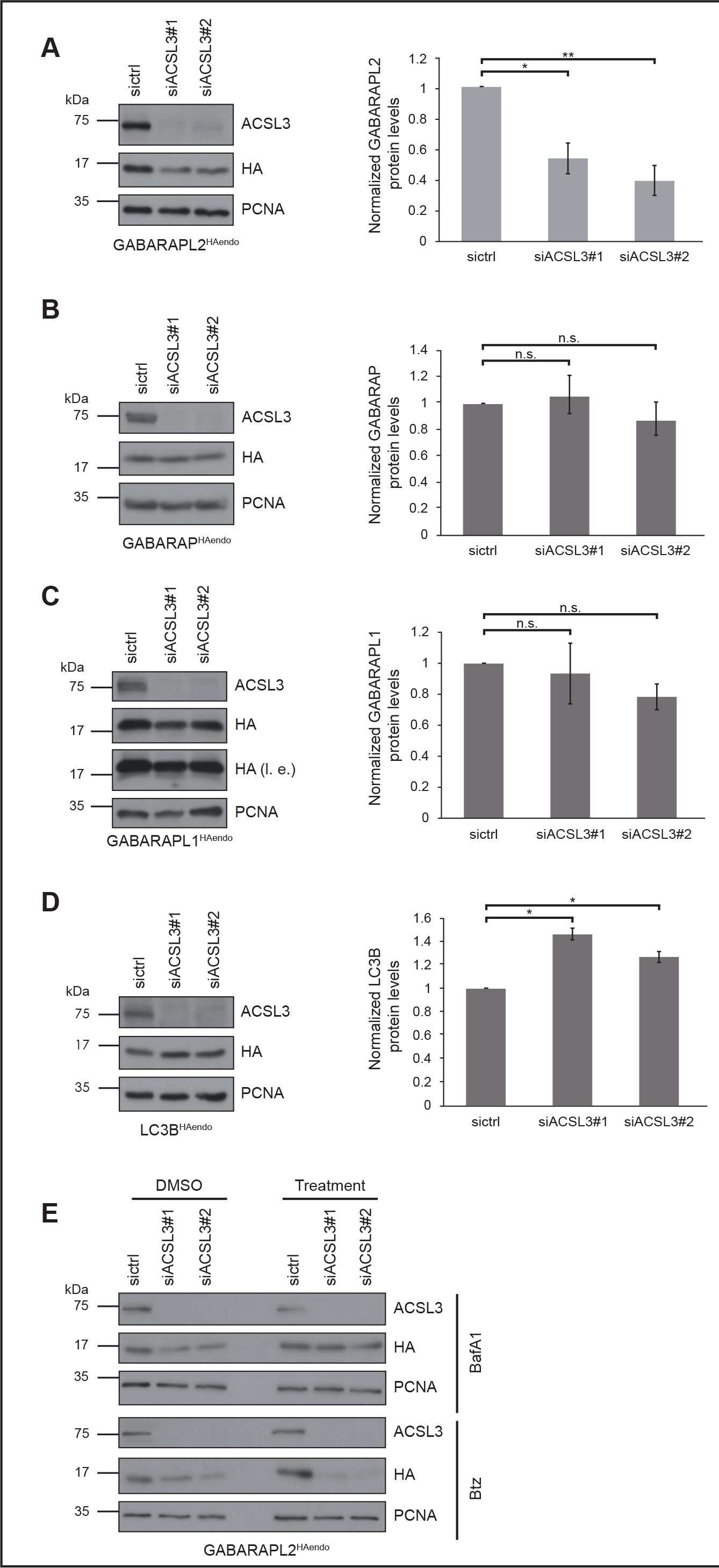
Stabilization of GABARAPL2 through ACSL3. ***A-D***, GABARAPL2_endoHA_ (***A***), GABARAP_endoHA_ (***B***), GABARAPL1_endoHA_ (***C***) and LC3B_endoHA_ (***D***) cells were reversely transfected with two different ACSL3 siRNAs. Lysates were analyzed by immunoblotting with indicated antibodies. Data represent mean ±SEM. Statistical analysis (n = 4) of the HA/PCNA ratio was performed using Student’s t-test (*p<0.005, **p<0.001). ***E,*** GABARAPL2_endoHA_ cells reversely transfected with siRNAs targeting ACSL3 for 72 hrs were treated with BafA1 or Btz and analyzed by immunoblotting.

Thus, these results indicate that ACSL3 is not degraded by autophagy but rather serves as a specific stabilizing factor of GABARAPL2 at the ER.

### ACSL3 anchors UBA5 to the ER membrane

To better understand the biological significance of the GABARAPL2-ACSL3 interaction, we turned our attention to known GABARAPL2 binding proteins and in particular to ubiquitin-like modifier activating enzyme 5 (UBA5) (32) which was recently shown to be recruited to the ER membrane in a GABARAPL2-dependent manner (33). Using HA- and myc-IPs, we confirmed the GABARAPL2-UBA5 interaction in GABARAPL2_endoHA_ cells transfected with myc-UBA5 (Figure 4A, Supplementary Figure S3C). Since ACSL3 binds GABARAPL2 at the ER membrane, we investigated whether ACSL3 also associates with UBA5. To this end, we generated HeLa cells stably overexpressing C-terminally HA-tagged ACSL3 and transiently transfected these and parental HeLa cells with myc-UBA5. Following differential treatment with oleic acid, cells were lysed and subjected to IP with anti-myc agarose. Intriguingly, we found that UBA5 associates with ACSL3 independent of its activity during LD formation (Figure 4B). Next, we examined whether ACSL3, GABARAPL2 and UBA5 form a ternary complex. Therefore, GABARAPL2_endoHA_/ACSL3_endoNeonGreen_ cells were treated with oleic acid or EtOH as control, followed by anti-UBA5 and anti-HA immunolabeling. Consistent with our binding assays, confocal microscopy showed colocalization of all three proteins irrespective of the treatment condition (Figure 4C). Overall, these results suggest that ACSL3, GABARAPL2 and UBA5 form a complex at the ER membrane independent of ACSL3’s activity in response to induction of LD formation.

### ACSL3 regulates ufmylation pathway components

Since we found that ACSL3 stabilizes GABARAPL2, we investigated whether ACSL3 depletion has similar effects on UBA5 protein abundance. For this purpose, GABARAPL2_endoHA_ cells were transfected with siACSL3 or sictrl and grown in the absence or presence of BafA1 or Btz. Indeed, we observed that protein levels of UBA5 decreased upon ACSL3 depletion and were restored by blockage of autophagy (Figure 5A). While depletion of GABARAPL2 had no effects on UBA5 protein levels (Supplementary Figure S3E). This supports the notion that UBA5 and GABARAPL2 form a functional unit which is regulated by ACSL3. UBA5 is part of the conjugation system, termed ufmylation, that covalently attaches the ubiquitin-like protein ubiquitin fold modifier 1 (UFM1) to target proteins through an E1-E2-E3 multienzyme cascade. The E1-like enzyme UBA5 activates UFM1 by forming a thioester bond between its active site and the exposed C-terminal glycine of UFM1 (32). The UFM1-conjugating enzyme 1 (UFC1) then transfers UFM1 from UBA5 to the UFM1-protein ligase 1 (UFL1) which mediates the attachment to target proteins (32, 34). The ER-membrane bound protein DDRGK1 anchors UFL1 to the ER membrane (35) and is besides RPL26 (36) and ASC1 (37) one of the few known ufmylation targets (34). While the consequences of ufmylation remains poorly understood at the mechanistic level, the UFM1 conjugation pathway has been linked to the ER stress response (38, 39), erythrocyte differentiation (40, 41), cellular homeostasis (42) and breast cancer progression (37). Since the stability of UBA5 and its ER-recruiting factor GABARAPL2 was controlled by ACSL3, we probed whether it also regulates the abundance of the other proteins in the ufmylation cascade. Knockdown experiments revealed that the protein levels of UFM1, UFL1 and DDRGK1 were likewise decreased upon ACSL3 depletion while the abundance of UFC1 was unaffected. Notably, UFC1 is the only ufmylation component that is not localized at the ER membrane. The observation that the protein levels of UFM1, UFL1 and DDRGK1 were not restored by blockage of autophagy or blockage of the proteasome (Figure 5B) indicates that these ufmylation factors are regulated at the transcriptional level. Together, our findings suggest that ACSL3 not only anchors UBA5 but might act as novel regulator of the ufmylation cascade.

**Figure 4.**
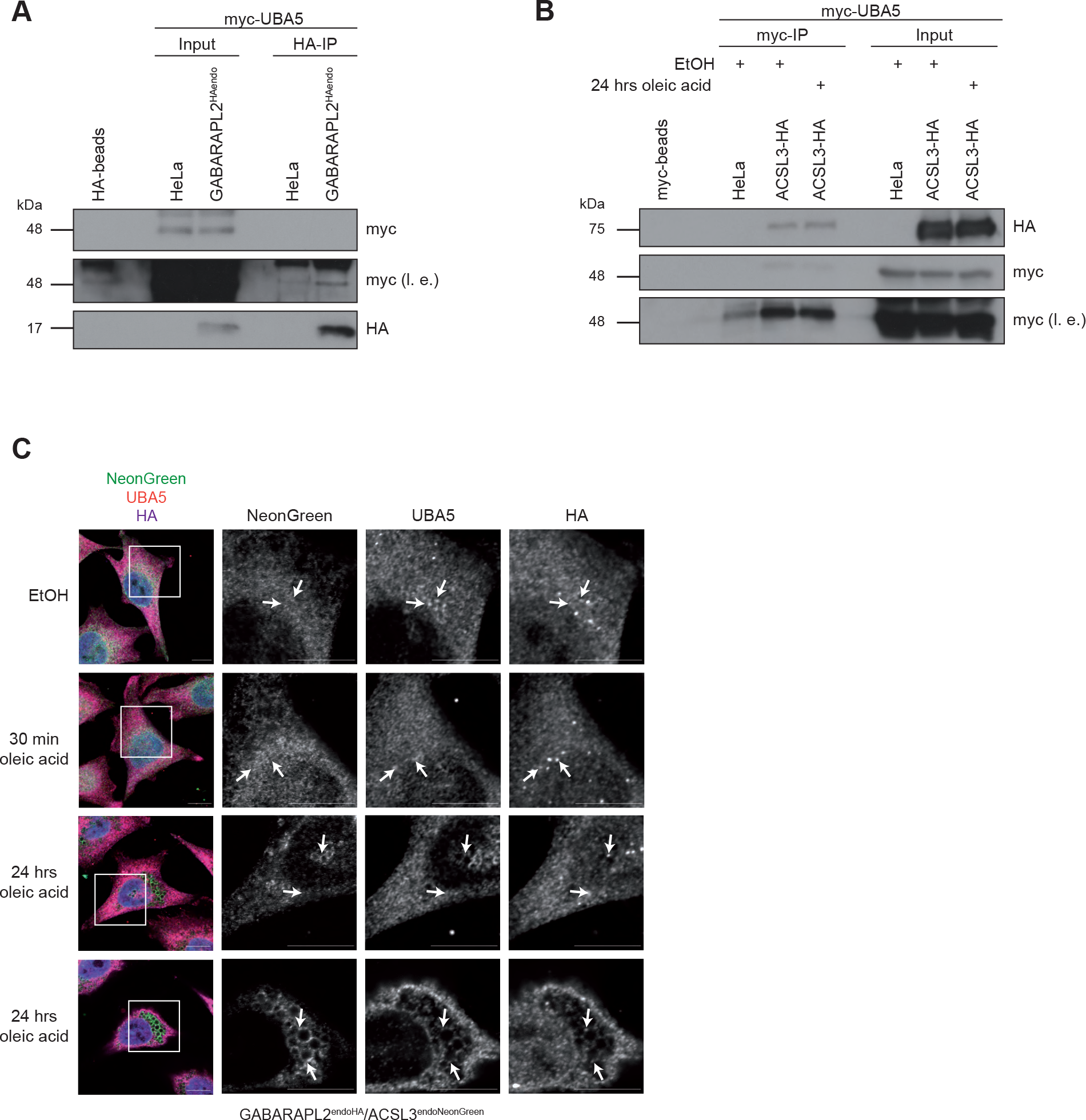
UBA5 binds to ACSL3 and GABARAPL2. ***A,*** Immunoblot analysis of anti-HA immunoprecipitates from lysates derived from parental HeLa and GABARAPL2_endoHA_ cells transiently transfected for 48 hrs with myc-UBA5. ***B,*** Parental HeLa and GABARAPL2_endoHA_ cells transfected with myc-UBA5 were treated with oleic acid or EtOH for 24 hrs prior to lysis, anti-myc immunoprecipitation and immunoblotting. ***C,*** GABARAPL2_endoHA_/ACSL3_endoNeonGreen_ cells were treated with oleic acid for 30 min or 24 hrs followed by fixation and immunolabeling with anti-UBA5 and -HA antibodies. Scale bar: 10 µm. Arrowheads indicate colocalization events between UBA5, HA and NeonGreen. Two confocal planes are shown for the 24 hrs time point.

**Figure 5.**
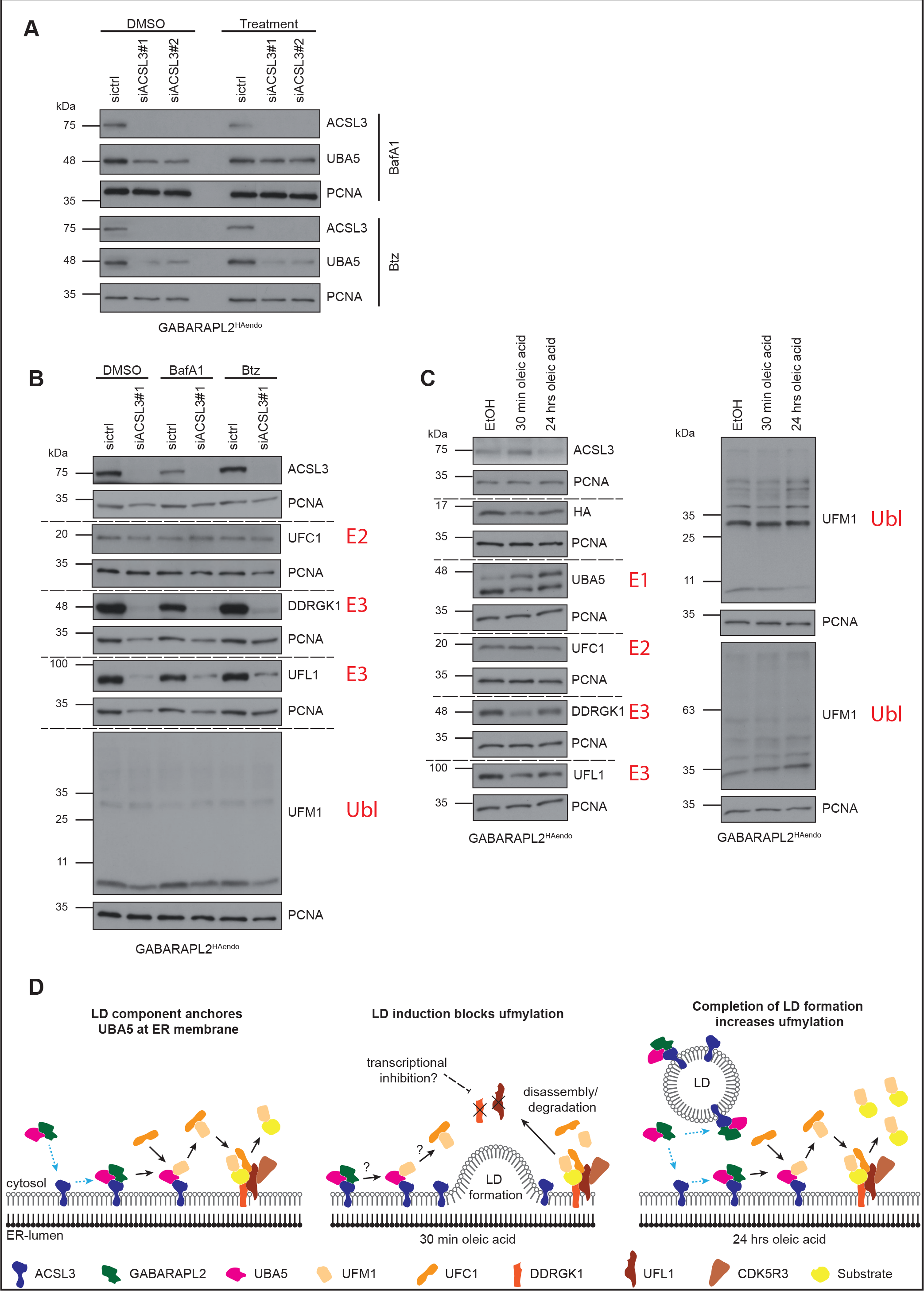
Influences of ACSL3 on the ufmylation pathway. ***A,B,*** GABARAPL2_endoHA_ cells were transfected with ACSL3 siRNAs and treated with Btz or BafA1 followed by lysis and immunoblot analysis using indicated antibodies. ***C,*** GABARAPL2_endoHA_ cells were treated with oleic acid for 30 min or 24 hrs or with EtOH for 24 hrs prior to lysis and immunoblotting with indicated antibodies. ***D,*** Working model summarizing the impact of ACSL3 and LD biogenesis on the ufmylation pathway. Upon recruitment via GABARAPL2, UBA5 is anchored at the ER membrane by ACSL3. LD induction through oleic acid blocks ufmylation through degradation-mediated disassembly of the UFM1 E3 enzyme complex. Completion of LD formation leads to reassemble of the E3 complex and increased ufmylation. Dotted blue arrows indicate ER-recruitment, black arrows indicate ufmylation cascade.

### LDs regulate UFM1 conjugation

These findings raise the question whether LD biogenesis and ufmylation are functionally coupled. To test this hypothesis, we monitored the ufmylation pathway in response to induction and completion of LD formation in GABARAPL2_endoHA_ cells grown in the absence and presence of oleic acid for 30 min and 24 hrs, respectively. While UBA5 levels drastically increased after 30 min and 24 hrs oleic acid treatment, there was no effect on UFC1 (Figure 5C, left panel). In contrast, the protein levels of DDRGK1 and UFL1 firstly decreased after 30 min incubation with oleic acid but after 24 hrs when LDs were formed DDRGK1 and UFL1 levels were partially restored (Figure 5C, left panel). Consistent with a restoration of the UFM1 conjugation pathway upon LD completion, we detected less unconjugated UFM1 (∼9kDa) and more conjugated UFM1 (∼35kDa) in response to 24 hrs oleic acid treatment (Figure 5C, right panel). Together, these results indicate that the ufmylation cascade is differentially regulated during induction and completion of LD and that the ACSL3-GABARAPL2-UBA5 axis plays an important part in this regulation.

## Discussion

In this study, we identified the ER-resident transmembrane protein ACSL3 as novel binding partner of GABARAPL2 and UBA5 using a CRISPR/Cas9 generated GABARAPL2_endoHA_ cell line. Furthermore, we provide evidences for the regulation of ufmylation through ACSL3 and LD biogenesis.

In our interactome screen with endogenously tagged GABARAPL2 we found ACSL3, which we confirmed as GABARAPL2 interactor by immunoprecipitation and confocal microscopy. Typically, interaction between GABARAPs or LC3s and their binding partners involves an ATG8 family-interacting motif (AIM; also known as LC3-interacting region (LIR)) in the hATG8 interactors and the LIR-docking site (LDS) in LC3 or GABARAP proteins (43–45). Amino acid sequence analysis of ACSL3 revealed four potential LIRs (LIR1: 65-71, LIR2: 135-140, LIR3: 589-594, LIR4: 643-648). While LIR2 is localized within the AMP-binding domain of ACSL3 and therefore is unlikely accessible, one (or all) of the other three ACSL3 LIR candidates may in principle mediate the binding to GABARAPL2. However, in addition to the LIR/LDS pairing Marshall and colleagues recently reported an alternative hATG8 interaction modus in which binding partners employ a ubiquitin-interacting motif (UIM) to bind to an UIM-docking site (UDS) in LC3 and GABARAP proteins (46, 47). By sequence inspection we found one potential UIM in ACSL3 (663-670). Given that the results from our colocalization studies and binding assays points to a possible complex formation of GABARAPL2, ACSL3 and UBA5 and that the LDS of GABARAPL2 is likely to be occupied by the atypical LIR of UBA5 (33, 48), it is highly plausible that GABARAPL2 and ACSL3 interact in a UIM/UDS-dependent manner.

GABARAP proteins were shown to mediate ER recruitment of UBA5 to bring it in close proximity to the membrane bound UFM1 E3 enzyme complex composed of UFL1, DDRGK1 and CDK5R3, thereby facilitating ufmylation (33). However, since GABARAPs are not known to be conjugated to PE at the ER, the molecular basis of this recruitment process was not clear. Here, we provided evidence that ACSL3 function to anchor UBA5 at the ER membrane. Given that UBA5 employs an atypical LIR to bind both GABARAPL2 and UFM1 and that the latter is able to outcompete GABARAPL2 binding of UBA5 *in vitro* (48), it is tempting to speculate that GABARAPL2 interacts with UBA5 until UFM1 conjugation is triggered. In this scenario, GABARAPL2 is only a recruiting factor that hands UBA5 over to ACSL3 (Figure 5D). However, the binding mode of ACSL3 and UBA5 remains to be explored.

While targets of ufmylation are still largely unknown, two of the three known UFM1-modified proteins are linked to the ER. Firstly, UFM1 conjugation of DDRGK1 is essential for the stabilization of the serine/threonine-protein kinase/endoribonuclease IRE1 (inositol-requiring enzyme 1) (37, 49). Secondly, it was shown that RPL26 (60S ribosomal protein L26) is exclusively ufmylated and de-ufmylated at the ER membrane (36). Overall, emerging evidence points to a role of the UFM1 conjugation system as regulator of ER homeostasis, ER stress response and ER remodeling. Disruption of protein folding and accumulation of unfolded proteins in the ER are hallmarks of ER stress which leads to the induction of the unfolded protein response (UPR) via one of these three key factors: IRE1, PKR-like ER protein kinase (PERK) or activating transcription factor 6 (ATF6). Protein degradation, reduction of protein synthesis and enlargement of the ER capacity are part of the UPR (50). In different cell lines and animal models, it was reported that ufmylation is upregulated via IRE1 or PERK upon ER stress, while depletion of ufmylation components induce the UPR (38, 39, 42, 51, 52). Upon re-established ER homeostasis, ufmylation coordinates the elimination of extended ER membranes through ER-phagy (53, 54).

In our present study, we identified LD formation stimulated by oleic acid treatment as novel regulator of ufmylation. LD biogenesis starts with lens formation, an accumulation of neutral lipids between the ER membrane leaflets until LDs eventually bud from the ER. The hydrophobic neutral lipid core of a LD is surrounded by a phospholipid monolayer with the origin of the outer ER membrane leaflet (55). ACSL3 was identified as LD associated protein and essential for LD biogenesis, expansion and maturation (27, 56). During initiation of LD biogenesis ACSL3 is translocated and concentrated to pre-LDs to drive LD expansion by mediating acyl-CoA synthesis. However, cells with enzymatically inactive ACSL3 are still able to form LDs, suggesting additional functions of ACSL3 in LD biogenesis (27, 57). Induction of LD formation induced by oleic acid resulted in an immediate (after 30 min) reduction of UFL1 and DDRGK1 protein levels and thus shut down of UFM1 conjugation (Fig 6A). Interestingly, depletion of ACSL3 led to a similar phenotype with regard to these two ufmylation components. Together, these results suggest that ACSL3 regulates DDRGK1 and UFL1 protein levels and therefore ufmylation. The observation that inhibition of proteasomal or lysosomal degradation only partly rescued this phenotype suggests that the ufmylation machinery is probably downregulated at the transcriptional level. To what extend this involves one of the three UPR factors IRE1, PERK or ATF6 remains to be examined. Considering that ER-phagy is blocked by inhibition of the interaction between DDRGK1 and UFL1 (53), we hypothesize that LD biogenesis inhibits the remodeling of ER membranes by ER-phagy. Intriguingly, completion of LD formation (∼24 hrs) almost restored DDRGK1 and UFL1 protein levels, which in turn led to increased ufmylation (Figure 5D). Whether these UFM1 targets are linked to remodeling of ER membranes by re-established ER-phagy remains to be tested.

Collectively, these findings underline the potential of our CRISPR/Cas9 gene-edited cell lines to uncover novel cellular pathways involving hATG8 family members without the need of overexpression systems, thereby complementing the recently generated LC3 and GABARAP knockout cell lines (9). Together with the LC3C_endoHA_ cell line that we previously reported (20) this cellular resource circumvents the drawback of unspecific LC3 and GABARAP antibodies and hence will greatly facilitate the functional dissection of individual hATG8 proteins.

## Material and Methods

### Cell culture and treatments

HeLa cell lines were cultured in Dulbecco’s modified Eagle’s medium (DMEM) + GlutaMAX-I (Gibco) supplemented with 10 % fetal bovine serum (FBS) and 1mM sodium pyruvate (Gibco) and grown at 37° C and 5 % CO_2_. For SILAC mass spectrometry, cells were grown in lysine- and arginine-free DMEM (Gibco) supplemented with 10 % dialyzed FBS, 2 mM glutamine (Gibco), 1 mM sodium pyruvate (Gibco) and 146 mg/ml light (K0, Sigma) or heavy L-lysine (K8, Cambridge Isotope Laboratories) and 84 mg/ml light (R0, Sigma) or heavy L-arginine (R10, Cambridge Isotope Laboratories). SILAC labeled cells were counted after harvesting, mixed 1:1 and stored at -80° C. For selection Puromycin (2 µg/ml) or Blasticidine (4 µg/ml) was added to the growth medium. The following reagents were used for treatments: oleic acid (EMD Millipore, 4954, 600 µm in EtOH, 30 min or 24 hrs), Bafilomycin A1 (Biomol, Cay11038-1, 200 nM in DMSO, 2 hrs), Torin 1 (Tocris, 4247, 250 nM in DMSO, 2 hrs), Bortezomib (LC Labs B-1408, 1 µM in PBS, 8 hrs), ATG7 inhibitor (Takeda ML00792183, 1 µM, 24 hrs).

### Plasmids

attB flanked ORFs, generated by PCR were cloned into the Gateway entry vector pDONR233. ORFs from pDONR233 constructs were introduced into one of the following destination vectors using recombination cloning: pHAGE-CMV-C-FLAG-HA, pEZYmyc-HIS (Addgene, #18701) or pDEST-myc. Stable pHAGE-ACSL3-HA expressing cells were generate by lentiviral transduction followed selection with 2 µg/ml Puromycin. pEZY and pDEST constructs were used for transient expression in cells (see transfection).

### Genome editing

The N-terminal HA-tagged hATG8 cell lines were generated with homology PCR templates containing 87 bp of GABARAP/GABARAPL1/GABARAPL2/LC3B-5’UTR including the start codon followed by the Blasticidine resistance gene, P2A, HA and 92bp downstream of the start codon of the corresponding hATG8 gene. For the C-terminal ACSL3-NeonGreen cell line, we used a homology PCR template containing 75 bp of the last exon of ACSL3, the NeonGreen ORF (Allele Biotech), T2A and the Blasticidine resistance gene ending with 84 bp downstream of the last exon of ACSL3. sgRNAs for hATG8s and ACSL3, designed with the online design tool from the Broad Institute (https://portals.broadinstitute.org/gpp/public/analysis-tools/sgrna-design) were clone into BSbI digested px330 (Addgene #42230), a SpCas9 expressing plasmid (sgRNA: GABARAP: GGAGGATGAAGTTCGTGTAC, GABARAPL1: TGCGGTGCATCATGAAGTTC, GABARAPL2: CCATGAAGTGGATGTTCAAG, LC3B: AGATCCCTGCACCATGCCGT, ACSL3: AGAAAATAATTATTCTCTTC). HeLa cells were seeded in a 6-well plate and transfected with Lipofectamin 2000 according to the manufacture’s instructions with sgRNA and corresponding homology PRC template. After 48 hrs, cells were selected with 4 µg/ml Blasticidine and single cell selection in 96-well plates. Correct introduction of the tag was verified by PCR and sequencing.

### Antibodies

For immunoblotting the following primary antibodies were used at a concentration of 1:1000 in 5 % milk-TBS-T: HA (Cell Signaling 3724S), HA (Biolegend, 901501), PCNA (Santa Cruz, sc-7907), ACSL3 (Santa Cruz sc-166374), mNeonGreen (Chromotek, 32F6), UBA5 (Proteintech 12093-1-AP/Sigma HPA017235), UFC1 (Proteintech 15783-1-AP), DDRGK1 (Proteintech 21445-1-AP), UFL1 (Abcam ab226216), UFM1 (Abcam ab109305) or at a concentration of 1:100 in 5 % milk-TBS-T: anti-myc (Monoclonal Antibody Core Facility, Helmholtz Zentrum Munich, 9E1, rat IgG1), anti-myc (Monoclonal Antibody Core Facility, Helmholtz Zentrum Munich, 9E10, mouse IgG). As secondary antibodies we used horseradish peroxidase coupled anti-mouse (Promega W402B), anti-rabbit (Promega, W401B) antibodies at a concentration of 1:10 000 and anti-rat IgG1 (Monoclonal Antibody Core Facility, Helmholtz Zentrum Munich) antibody at a concentration of 1:100 in 1 % milk-TBS-T. The following primary antibodies were used for immunofluorescence in 0.1 % BSA-PBS: HA (Roche, 11867423001, 1:50), p62 (BD, 610832, 1:500), LAMP1 (DSHB, H4A3, 1:50), LC3 (MBL, PM036, 1:500), HCS LipidTOX™ Red Phospholipidosis Detection Reagent (Thermo Scientific, H34351, 1:1000), HCS LipidTOX™ Deep Red Neutral Lipid Stain (Thermo Scientific, H34477, 1:500), UBA5 (Proteintech, 12093-1-AP, 1:250), Calnexin (Stressgen, SPA-860, 1:100), SEC13 (Novus, AF9055-100, 1:300). The following fluorophore conjugated secondary antibodies from Thermo Fisher were use at a concentration of 1:1000 in 0.1 % BSA-PBS: anti-mouse IgG Alexa Fluor 488 (A-11001), anti-rabbit IgG Alexa Fluor 488 (A-11008), anti-rabbit IgG Alexa Fluor 594 (A-11012) and anti-rat IgG Alexa Fluor 647 (A-21247).

### Transfection

For siRNA knockdowns, cells were reversely transfected with Lipofectamine RNAiMax (Thermo Fisher Scientific) according to the manufacturer’s guidance with 30 nM of the following siRNAs from Dharmacon/Horizon Discovery and harvested 72 hrs after transfection: sicrtl UGGUUUACAUGUUUUCCUA, siACSL3#1 UAACUGAACUAGCUCGAAA, siACSL3#2:

GCAGUAAUCAUGUACACAA, siGABARAP GGUCAGUUCUACUUCUUGA, siGABARAPL1 GAAGAAAUAUCCGGACAGG, siGABARAPL2 GCUCAGUUCAUGUGGAUCA, siLC3B GUAGAAGAUGUCCGACUUA. Plasmids were transiently transfected with Lipofectamine 2000 (Thermo Fisher Scientific) according to the instruction of the manufacturer or with 10 mM PEI (Polyethylenimine) and cells were collected after 48 hrs.

### Immunoblotting

Cell were lysed in RIPA (50 mM Tris-HCl [pH 7.4], 150 mM NaCl, 0.5% sodium desoxycholate, 1 % NP-40, 0.1 % SDS, 1x EDTA-free protease inhibitor (Roche), 1x phosphatase inhibitor (Roche)) for 30 min. After elimination of cell debris by centrifugation, proteins were diluted with 3x loading buffer (200 mM Tris-HCl [pH 6.8], 6 % SDS, 20 % Glycerol, 0.1 g/ml DTT, 0.1 mg Bromophenol blue) and boiled at 95°C. Proteins were size separated by SDS-PAGE with self-casted 8 %, 10 %, 12 % and 15 % gels followed by protein transfer onto nitrocellulose membranes (GE Healthcare Life Sciences, 0.45 µm). For better visibility of endogenous HA-hATG8s membranes were boiled for 5 min in PBS after protein transfer. Blots were blocked in TBS-T (20 mM Tris, 150 mM NaCl, 0.1% Tween-20) supplemented with 5 % low fat milk (Roth) for 1 hr. Primary antibodies were incubated overnight followed by several wash steps with TBS-T and incubation with secondary antibodies for 1 hr at room temperature. After repeated washing, immunoblots were analyzed with Western Lightning Plus ECL (Perkin Elmer).

### Immunofluorescence

All steps were carried out at room temperature. Cells growing on glass coverslips in 12-well plates were fixed with 4 % paraformaldehyde in PBS for 30 min followed by permeabilization with 0,1 % Trition-X-100 in PBS or 0,1 % Saponin in PBS for 30 min and 1 hr blocking in 1% BSA-PBS. First and secondary antibody incubation was done sequentially for 1 hr at room temperature in 0.1 % BSA-PBS followed by mounting of the coverslips with ProlongGold Antifade with Dapi (Thermo Fisher). In between each step, cells were washed several times with PBS. Cells were imaged with a LSM 800 Carl Zeiss microscope using 63x oil-immersion objective and ZEN blue edition software and analyses with ImageJ (version 1.52).

### Immunoprecipitation

Frozen cell pellets from 4×15 cm cell culture plates for mass spectrometry or 1×15 cm cell culture plate for immunoblotting were lysed in Glycerol buffer (20 mM Tris [pH 7.4], 150 mM NaCl, 5 mM EDTA, 0.5 % Triton-X-100, 10 % Glycerol, 1x protease inhibitor, 1x phosphatase inhibitor) for 30 min at 4° C with end-over-end rotation. Lysates were cleared from cell debris by centrifugation prior to adjustment of protein concentrations between the samples and overnight immunoprecipitation at 4° C with pre-equilibrated anti-HA-agarose (Sigma) or anti-c-myc-agarose (Thermo fisher). Agarose beads were washed five times with Glycerol buffer followed by elution of proteins with 3x loading buffer and boiling of the samples at 95° C. Samples were then analyzed by SDS-PAGE (self-casted or BioRad’s 4-20 % gels) followed by immunoblotting or in-gel tryptic digestion.

### Mass spectrometry

SDS-PAGE gel lines were cut in 12 equal size bands, further chopped in smaller pieces and placed in 96 well plates (one band per well). Gel pieces were washed with 50 mM ammonium bicarbonate (ABC)/50 % EtOH buffer followed by dehydration with EtOH, reduction of proteins with 10 mM DTT in 50 mM ABC at 56° C for 1 hr and alkylation of proteins with 55 mM iodacetamide in 50 mM ABC at room temperature for 45 min. Prior to overnight trypsin-digest (12 ng/ul trypsin in 50 mM ABC, Promega) at 37° C, gel pieces were washed and dehydrated as before. Peptide were extracted from gel pieces with 30 % acetonitrile/3 % trifluoroacetic acid (TFA), 70 % acetonitrile and finally 100 % acetonitrile followed by desalting on custom-made C18-stage tips. Using an Easy-nLC1200 liquid chromatography (Thermo Scientific), peptides were loaded onto 75 µm x 15 cm fused silica capillaries (New Objective) packed with C18AQ resin (Reprosil-Pur 120, 1.9 µm, Dr. Maisch HPLC). Peptide mixtures were separated using a gradient of 5%–33% acetonitrile in 0.1% acetic acid over 75 min and detected on an Q Exactive HF mass spectrometer (Thermo Scientific). Dynamic exclusion was enabled for 30 s and singly charged species or species for which a charge could not be assigned were rejected. MS data were processed with MaxQuant (version 1.6.0.1) and analyzed with Perseus (version 1.5.8.4, http://www.coxdocs.org/doku.php?id=perseus:start). IP experiments from GABARAPL2_endoHA_ and control parental HeLa cells were performed in duplicates and triplicates, respectively. Matches to common contaminants, reverse identifications and identifications based only on site-specific modifications were removed prior to further analysis. Log2 heavy/light ratios were calculated. A threshold based on a log2 fold change of greater than 1.5-fold or less than -1.5-fold was chosen so as to focus the data analysis on a smaller set of proteins with the largest alterations in abundance. Additional requirements were at least two MS counts, unique peptides and razor peptides as well as absence in IPs from parental HeLa control cells. For functional annotations, the platform DAVID (https://david.ncifcrf.gov/) was used.

### Data availability

The mass spectrometry proteomics data have been deposited to the ProteomeXchange Consortium (http://proteomecentral.proteomexchange.org) via the PRIDE partner repository with the dataset identifier PXD016734.

### Statistical analysis

Quantification and statistical analysis of western blots were done with imageJ and Phyton (version 3.7). Statistical significance was calculated with Student’s t test and data represent ± SEM (standard error of the mean). Statistical analysis of MS data was done with Perseus.

## Acknowledgement

We would like to thank Georg Werner and all members of the Behrends lab for reagents, advice and critical discussion. This work was supported by the Deutsche Forschungsgemeinschaft (German Research Foundation) within the framework of the Munich Cluster for Systems Neurology (EXC2145 SyNergy), the Collaborative Research Center (CRC1177), and the project grant BE 4685/2-1.

## Author contribution

FE performed all experiments. MK provided advice for CRISPR/Cas9 tagging strategy. FE and CB conceived the study and wrote the manuscript.

## Supplementary Figure Legends

**Supplementary Figure S1.**
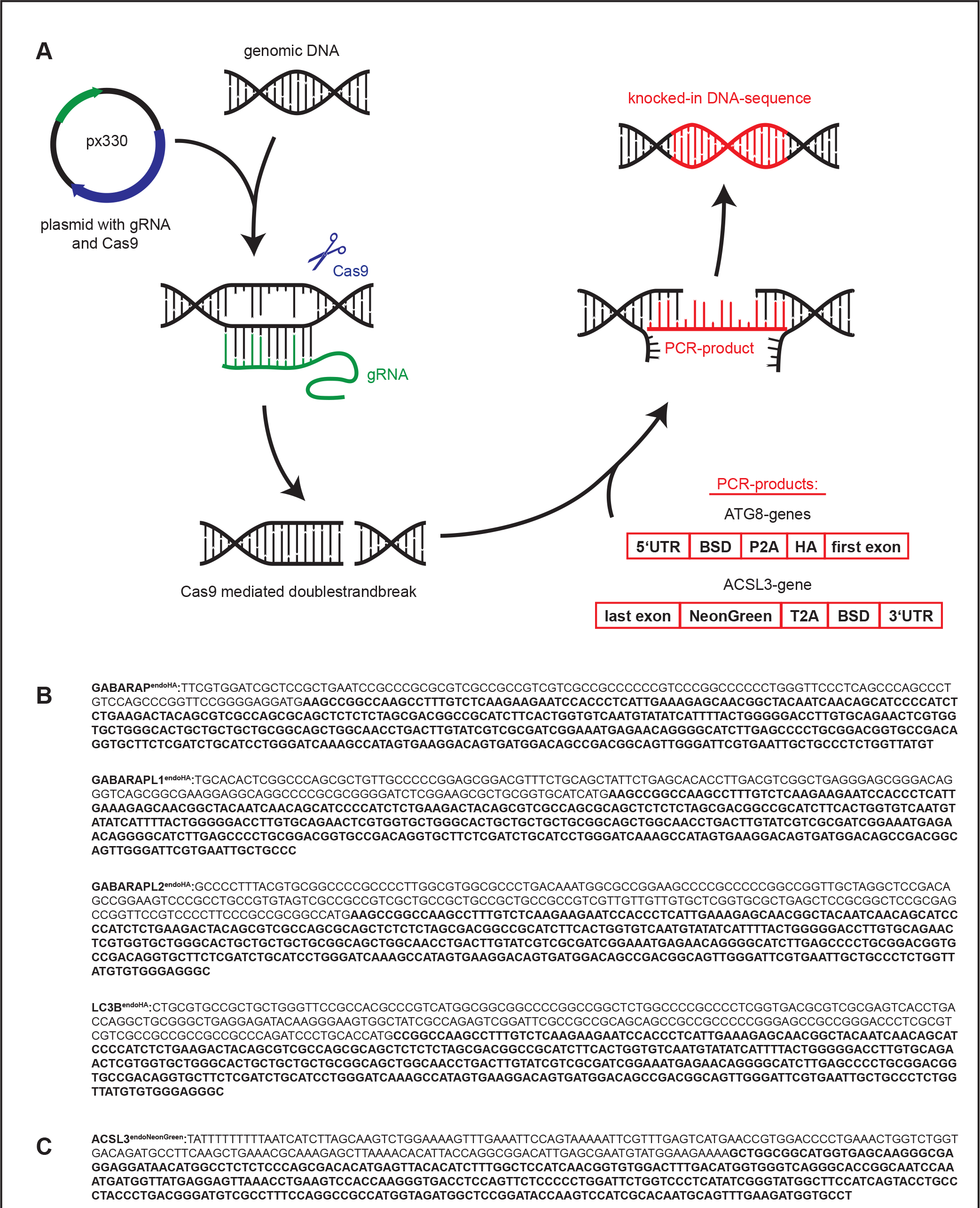
Endogenous epitope tagging of hATG8 and ACSL3 genes. ***A***, Experimental CRISPR/Cas9 workflow. ***B***,***C,*** Sequence data from PCR products of the tagged GABARAP_endoHA_, GABARAPL1_endoHA_, GABARAPL2_endoHA_, LC3B_endoHA_ cell lines (***B***) and the GABARAPL2_endoHA_/ACSL3_endoNeonGreen_ cell line (***C***). Sequence data from PCR products of the. Introduced CRISPR sequences are indicated in bold.

**Supplementary Figure S2.**
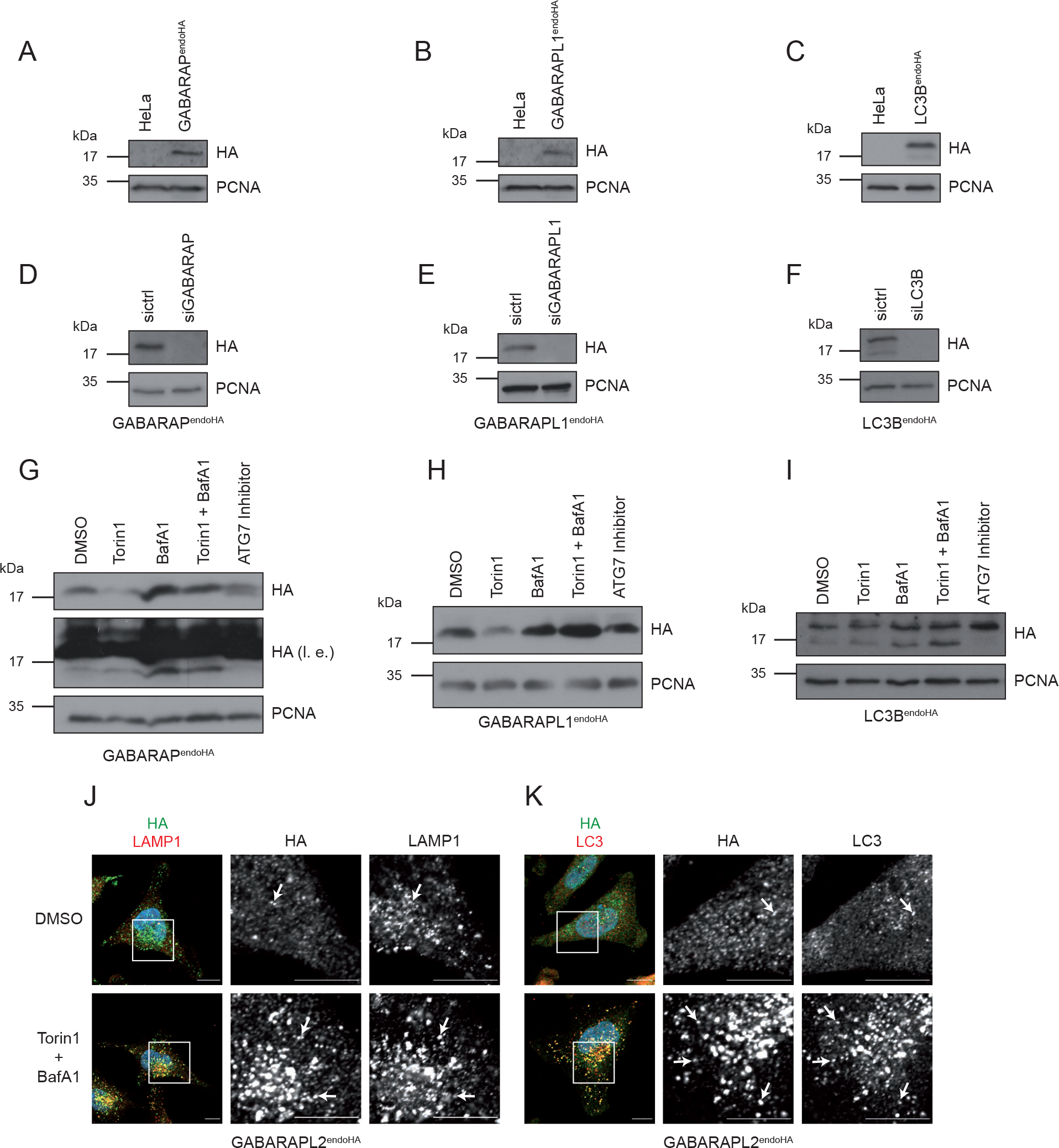
Validation of endogenously HA-tagged hATG8 proteins. ***A-C,*** GABARAP_endoHA_ (***A***), GABARAPL1_endoHA_ (***B***), LC3B_endoHA_ (***C***) and parental HeLa (***A-C***) cells were lysed followed by immunoblotting and analysis with indicated antibodies. ***D-F***, GABARAP_endoHA_ (***D***), GABARAPL1_endoHA_ (***E***), LC3B_endoHA_ (***F***) cell lines were reversely transfected with indicated siRNAs prior to immunoblot analysis. ***G-I,*** GABARAP_endoHA_ (***G***), GABARAPL1_endoHA_ (***H***), LC3B_endoHA_ (***I***) were treated as indicated followed by lysis and immunoblotting. ***J,K,*** GABARAPL2_endoHA_ cells treated with indicated inhibitors were immunolabeled with anti-LAMP1 (***K***) or anti-LC3 (***L***) antibody. Scale bar: 10 µm. Arrowheads indicate colocalization events.

**Supplementary Figure S3.**
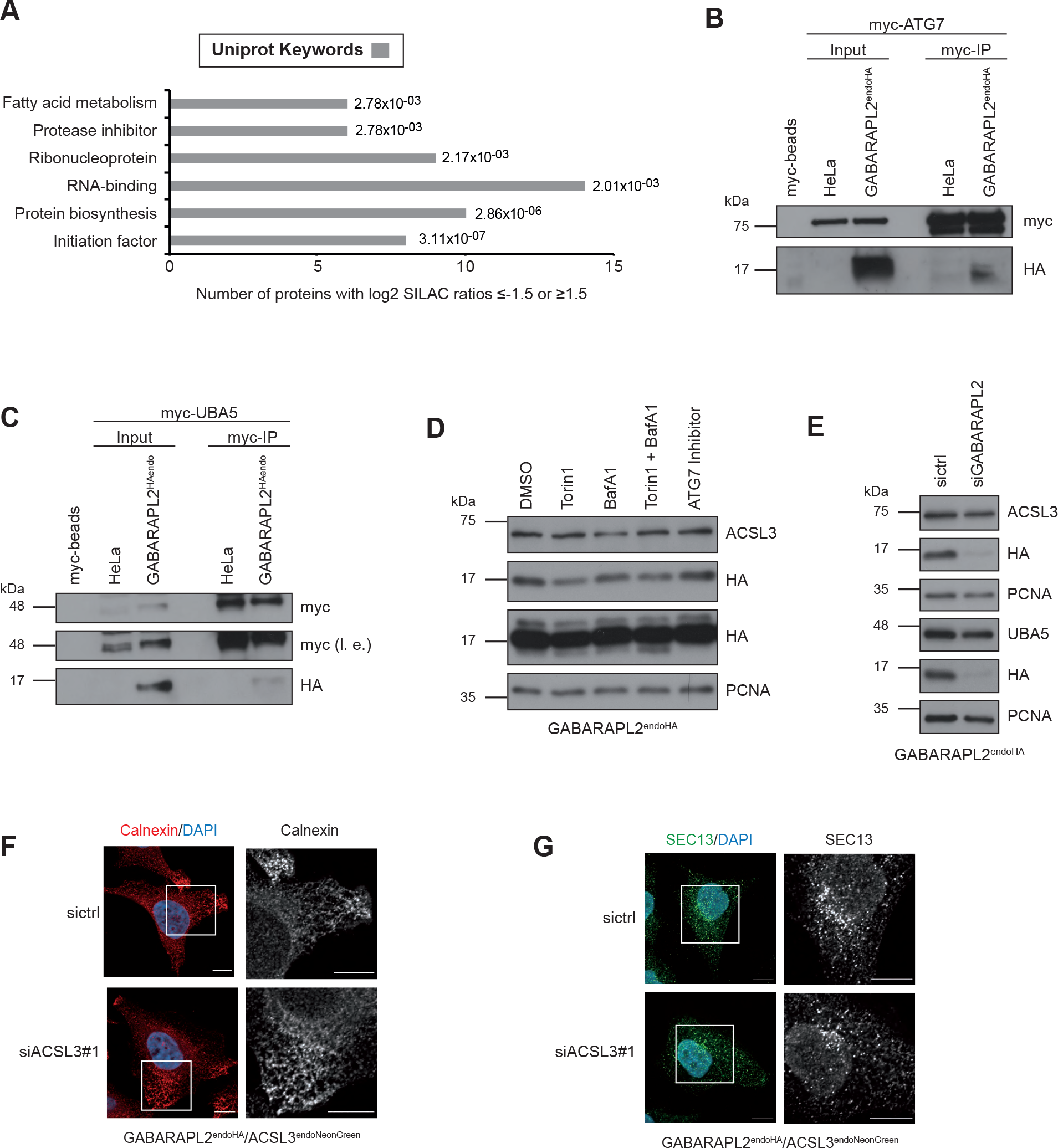
Analysis of GABARAPL2-interacting proteins. ***A,*** Annotation enrichment analysis of candidate GABARAPL2-interacting proteins with log2 SILAC H/L ratios ≥1.5 or ≤-1.5. The bar graphs show significantly overrepresented UniProt keywords. ***B,C,*** Immunoblot analysis of anti-myc immunoprecipitates from lysates derived from parental HeLa and GABARAPL2_endoHA_ cells transiently transfected for 48 hrs with myc-tagged ATG7 (***B***) or UBA5 (***C***). Stability and knockdown of ACSL3. ***D***, GABARAPL2_endoHA_ cells were treated as indicated and subjected to lysis and analyzed with immunoblotting and anti-ACSL3 antibody. ***E,*** Reversely transfected GABARAPL2_endoHA_ cells with non-targeting (sicrtl) or GABARAPL2 siRNA were lysed followed by immunoblotting and analysis with indicated antibodies. ***F,G,*** GABARAPL2_endoHA_/ACSL3_endoNeonGreen_ cells were transfected with indicated siRNAs prior to immunolabeling with Calnexin or SEC13. Scale bar: 10 µm.

